# Partitioning of semisynthetic lipidated N-Ras in lipid raft nanodomains determined by FRET to lipid domain markers

**DOI:** 10.1101/221382

**Authors:** Anna K. Shishina, Elizaveta A. Kovrigina, Azamat R. Galiakhmetov, Rajendra Rathore, Evgenii L. Kovrigin

## Abstract

Cellular membranes are heterogeneous planar lipid bilayers displaying lateral phase separation with the nanometer-scale liquid-ordered phase (aka “lipid rafts” or L_o_) surrounded by the liquid-disordered phase (L_d_). Many membrane-associated proteins were found to stably integrate in the rafts, which is critical for their biological function. Isoforms H and N of Ras GTPase possess a unique ability to switch their lipid domain preference depending on the type of bound guanine nucleotide (GDP or GTP). This behavior, however, has never been reproduced *in vitro* in model bilayers with recombinant proteins, and therefore has been attributed to action of other proteins binding Ras at the membrane surface. In this paper, we report the observation of the nucleotide-dependent switch of lipid domain preferences of the semisynthetic lipidated N-Ras in raft lipid vesicles *in the absence of other proteins*. To detect segregation of Ras molecules in raft and disordered lipid domains, we measured Förster Resonance Energy Transfer (FRET) between the donor fluorophore, mant, attached to the protein-bound guanine nucleotides, and the acceptor, rhodamine-conjugated lipid, localized to the liquid-disordered domains. We demonstrated that N-Ras preferentially populated raft domains when bound to mant-GDP, while losing preference for rafts when it was associated with a GTP mimic, mant-GppNHp. At the same time, the isolated lipidated C-terminal peptide of N-Ras was found localized outside of the liquid-ordered rafts, most likely—in the bulk disordered lipid.

## Introduction

Lipid rafts, the nanoscale lipid domains, in a plasma membrane of living cells play a crucial organizing role in cellular signaling and regulatory cascades^*1–6*^. Micrometer-sized lipid domains with a liquid-crystal-like order are easily observable by optical fluorescence microscopy in model membranes made of heterogeneous lipid mixtures^*4, 7–10*^. However, their cellular counterparts are expected to be much smaller, nanometer-sized, making them only resolvable by electron and atomic force microscopy techniques^11-14^. In a cell, many membrane proteins permanently reside in raft membrane domains, which is essential for their function^*5, 15–19*^. Ras, a small monomeric GTPase, provides an intriguing example of a membrane protein that dynamically switches its nanodomain affinity upon transition between its active and inactive functional states (bound to GTP and GDP, respectively)^*20–23*^.

Ras is a small monomeric GTPase involved in regulation of cell growth, proliferation and differentiation^*24*^. Mutations in the Ras genes are observed in up to 25% of all human cancers, which makes Ras one of the major targets for cancer therapy^*25–28*^. Ras consists of a GTPase catalytic domain (G domain) binding guanine nucleotides and the C-terminal peptide anchored to the inner leaflet of plasma membrane through a posttranslational lipidation motif^*29–31*^. Membrane attachment is crucial to Ras function: most effector proteins can only be activated by Ras-GTP when it is attached to the membrane surface^*25, 32*^.

Ras proteins are represented by three Ras isoforms with a high degree of homology and nearly 90% sequence identity in the N-terminal GTPase domain^*33*^. The remaining C-terminal 22-23 amino acids, known as the hyper-variable region, have no sequence similarity except for the conserved CAAX motif necessary for membrane targeting^*34*^. The variability of the C-terminal sequences of the Ras isoforms leads to different processing patterns in the cell. All Ras isoforms are modified by attachment of a prenyl (farnesyl) chain at the extreme C-terminal cysteine. H-Ras and N-Ras additionally get two and one palmitoyl chains, respectively, while K-Ras4B features a polybasic domain as alternative membrane-anchoring mechanism^*29, 35*^. It was demonstrated that the membrane-targeting region is responsible for partitioning of proteins between membrane domains^*36*^.

Dynamic change in H-Ras localization from cholesterol-rich rafts to the disordered lipid domains upon activation (GTP binding) was first observed using density gradients and immuno-gold electron microscopy in native cellular membranes^*22, 37–40*^. Explanation of this behavior of H-Ras was proposed when scaffolding protein galectin-1 was found to associate with activated H-Ras nanoclusters in disordered lipid domains^*41*^. The K-Ras isoform was found residing in the disordered phase irrespective of its activation status (bound GDP or GTP)^*22, 38*^. The lipid domain preferences of N-Ras remain controversial as it was observed in a raft phase of COS-7 cell membranes when in the GDP form^*42*^, while Roy reported that N-Ras-GDP was localized in the disordered lipid phase of BHK cells and moved to raft domains upon GTP binding^*43*^. Experiments in model membranes recapitulated none of these findings: N-Ras was found concentrated at the raft/disordered domain boundary in model lipid bilayers irrespective of the bound nucleotide^*44–47*^. The cited reports characterized N-Ras behavior in very different membranes systems: from natural plasma membranes of BHK and COS-7 cells to synthetic lipid mixtures, which might be one of the causes of difference. The dynamic shift from one phase to another upon activation of N-Ras observed by Roy et al.^*43*^ could, potentially, be due to binding to yet-unidentified protein scaffolds (by analogy with H-Ras). In the present report, we make use of a full-length semi-synthetic lipidated N-Ras to demonstrate that it is capable of changing its nano-domain localization in model lipid membranes in nucleotide-dependent manner in the absence of any other proteins.

## Results

Our goal was to assess relative affinity of N-Ras lipoprotein to raft and disordered lipid domains in a model lipid system, and determine whether raft affinity of N-Ras is dependent on the nature of a bound nucleotide (hence, the biologically active/inactive protein conformation) in the absence of “helper” proteins. Because of the nanoscale dimensions of rafts, we relied on measurements of FRET between Ras-attached fluorophore and fluorescent lipid domain markers^*48–50*^. H-Ras localization was previously probed by FRET to lipid domain markers but those reports did not include N-Ras^*51, 52*^.

In the following subsections we

1. evaluated the model lipid bilayers to confirm that they form nanometer ordered domains mimicking size of cellular rafts,
2. detected non-raft localization of the C-terminal lipidated peptide of N-Ras,
3. evaluated a hypothesis that the C-terminal peptide may be attracted to the raft boundary,
4. established lifetime-based detection of nanodomain localization, and
5. detected distinct nanodomain preferences of N-Ras in active and inactive states (bound to GTP mimic or GDP).

### 1. Lipid membrane mimic with nano-scale lipid domains

To create lipid bilayers that spontaneously forms nanometer-sized raft domains (approx. ranging from 4 to 15 nm), we followed Pathak and London^*50*^ and utilized a lipid mixture of sphingomyelin (SM), 1-palmitoyl-2-oleoyl-sn-glycero-3-phosphocholine (POPC), and cholesterol in the equimolar ratio (referred to in the following as the raft lipid mixture). A pure POPC lipid was used to make a homogeneous (non-raft) control bilayers. Confocal fluorescence microscopy of a supported lipid bilayer made of the raft lipid mix confirmed that these bilayers do not form non-physiological *micrometer-sized* rafts (Figure 1). We used the NBD-DPPE (green fluorescence) as a lipid raft marker and the Rhod-DOPE (red fluorescence) as a disordered domain marker to detect micrometer-sized domains. Figure 1 reveals the absence of significant areas of green and red color confirming that the size of rafts and disordered domains in this bilayer is on the order of or smaller than the optical resolution of the microscope, ca. 200 nm (compare to Crane and Tamm, 2007^*10*^).

**Figure 1.**
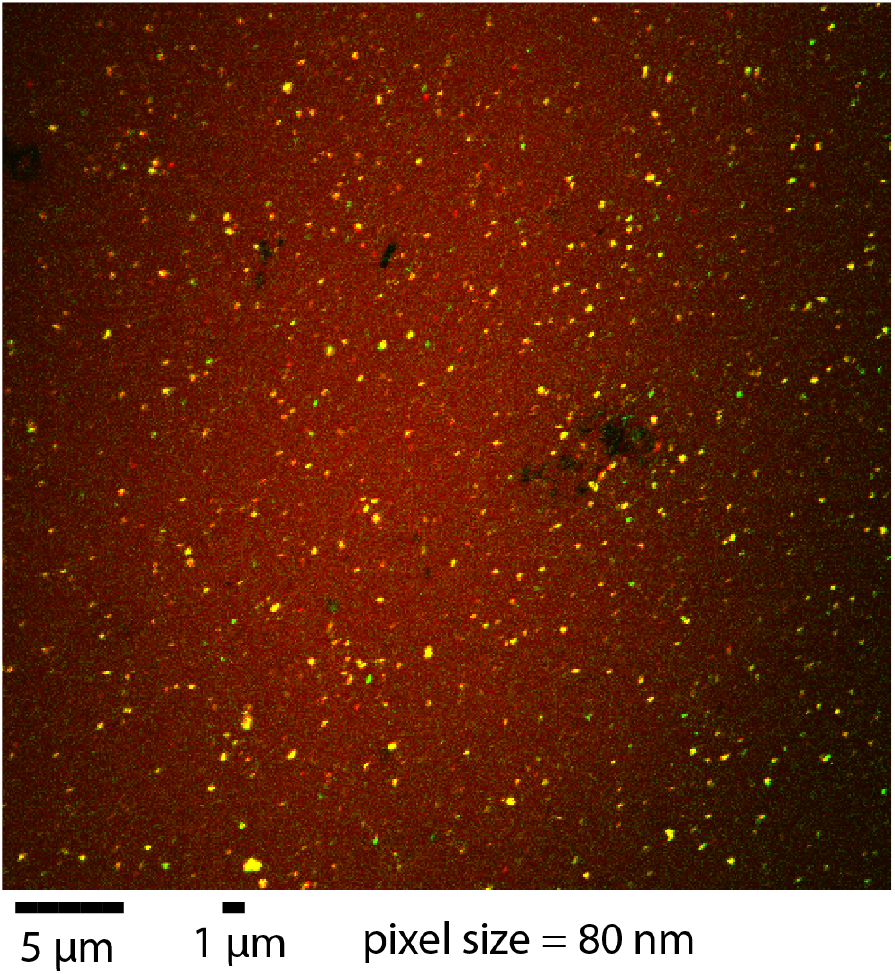
Overlay of images of NBD-DPPE fluorescence (green) and Rhod-DOPE fluorescence (red) in supported lipid bilayers made of the raft lipid mix (molar ratio: SM/POPC/Chol = 1:1:1). Bright yellow spots correspond to aggregated LUV that were not removed during the wash phase. Black areas (in the middle of the image) are, likely, due to defects on the glass surface.

To detect the presence of *nanoscopic* lipid rafts, we measured FRET between donors and acceptors segregated in raft or non-raft lipid domains^*48*^. The 1,6-diphenyl-1,3,5-hexatriene (DPH) served as a donor that is uniformly distributed in a lipid bilayer, while Rhod-DOPE was an acceptor that is preferentially excluded from the lipid rafts^*53–55*^. Therefore, in a raft lipid membrane, some fraction of DPH is segregated from Rhod-DOPE. Heating of LUV samples leads to melting of rafts and increased quenching of DPH fluorescence by Rhod-DOPE relatively to a homogeneous bilayer. The R_o_ of 36Å allows for sensitive detection of formation and melting of nanoscopic raft nanodomains^*50*^.

To demonstrate the nanoscale domain segregation in raft LUV, we recorded the fluorescence signal of DPH in the absence and the presence of Rhod-DOPE (F_o_ and F samples, respectively; DPH concentration held constant) as a function of temperature (Figure 2A). Quenching of the DPH signal by rhodamine manifested itself in a reduced ratio of fluorescence intensity of F to F_o_ sample. Due to difficulty with complete subtraction of the excitation light scattered by LUV, the absolute values of F to F_o_ intensity ratio on the Y axis contained both FRET and scattering contributions. However, the light scattering by LUV is relatively temperature-independent, therefore, the F/F_o_ variation with temperature reflects the relative change of FRET from DPH to Rhod-DOPE. Similarity of the overall shape of the profiles obtained upon heating and cooling confirmed reversibility of the measurement and relative photostability of the fluorophores. The raft LUV samples revealed a characteristic sigmoidal transition indicating relative segregation of acceptors from donors at low temperatures, and increased access of acceptors to donors upon heating due to melting of the lipid rafts (reduction in average size^*50*^) — schematically illustrated in Figure 2B. As anticipated, the homogeneous LUV control did not reveal dramatic changes in F/F_o_ upon heating.

**Figure 2.**
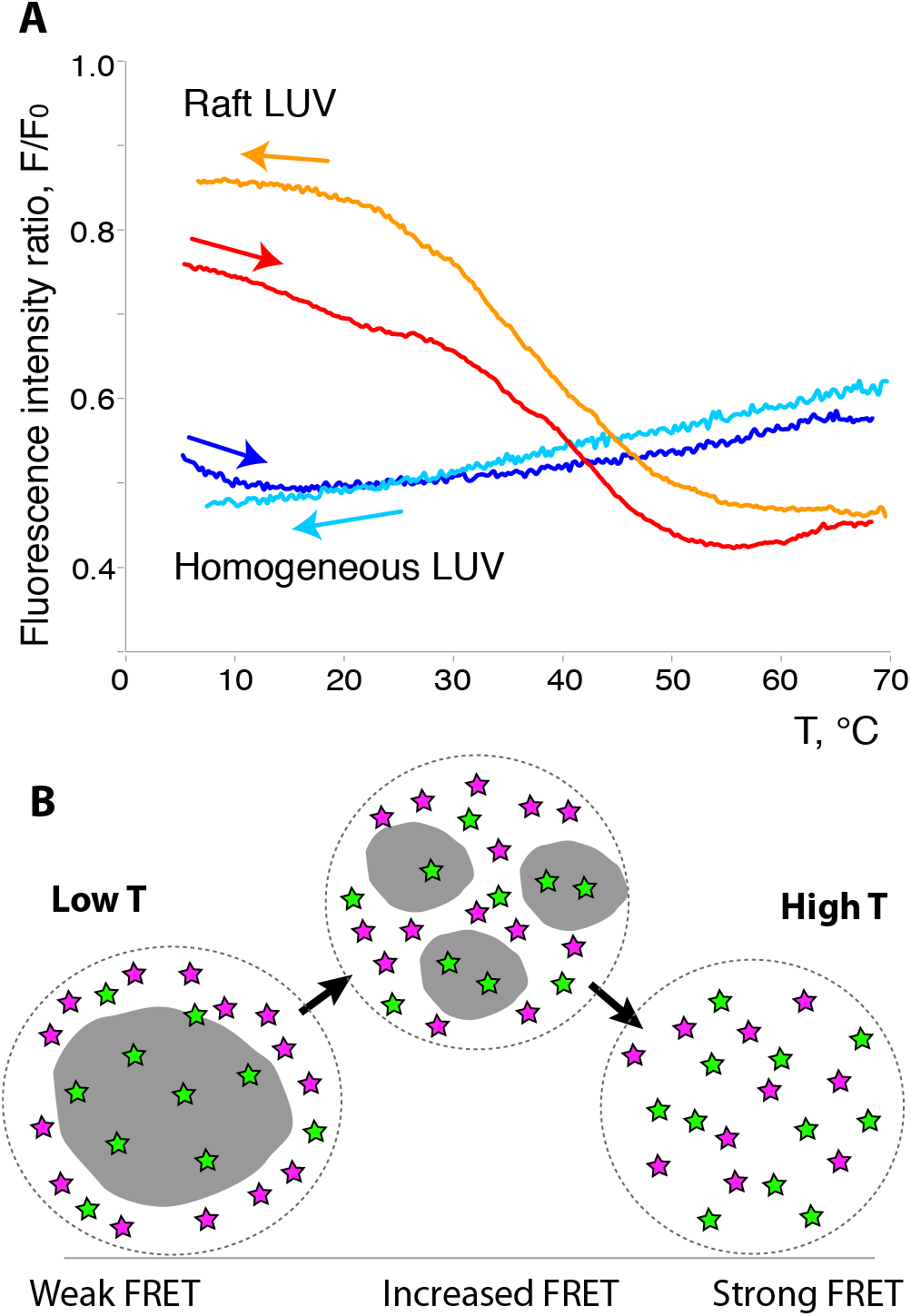
Presence of rafts in SM/POPC/cholesterol lipid bilayers detected by FRET between lipid domain markers. (A) Heating and cooling profiles of the homogeneous and raft LUV solutions with the DPH (0.1% mol) and Rhod-DOPE (2% mol) donor/acceptor pair at a scan rate of 0.5 ^*o*^C/min. Each curve is an average of two independent samples. Fluorescence intensity ratio, F/F_o_, is calculated using DPH emission of F and F_o_ samples, containing and lacking Rhod-DOPE, respectively. (B) A schematic drawing illustrating the increase of relative access of Rhod-DOPE acceptors (magenta stars) to the DPH donor molecules (green stars) upon heating. Gray area represents a lipid raft.

### 2. Preferential localization of N-Ras C-terminal lipopeptide

To determine a contribution of a lipidated C-terminus of N-Ras to the protein interactions with raft and non-raft lipid domains, we evaluated the preferential localization of N-Ras C-terminal lipopeptide in absence of the G domain. The N-methylanthranyl group (mant) was attached to the lipopeptide N-terminus to serve as a donor fluorophore. The mant is a small and relatively soluble fluorophore that was successfully utilized in a number of biochemical studies and demonstrated to induce minimal perturbations in biomolecular systems^*56–58*^. Spectral properties of mant are comparable to DPH, therefore, we expected a similar Förster radius and a similar sensitivity to the domain localization.

Heating and cooling profiles of F/F_o_ for mant-lipopeptide (Figure 3) revealed a pattern, which was opposite to the one observed for DPH in Figure 2A. Heating led to increasing F/F_o_ indicating reduction of the FRET efficiency at higher temperatures, while homogeneous lipid showed relatively constant F/F_o_ values. In an analogous system, Fastenberg et al. explained such increasing pattern by hypothesizing that donor is present in the same disordered phase as the acceptor^*59*^. In this scenario, melting of rafts upon heating leads to an increase in the disordered phase area and spreading of donor and acceptor fluorophores in the membrane plane, which results in reduction of FRET efficiency (greater intensity of F samples).

**Figure 3.**
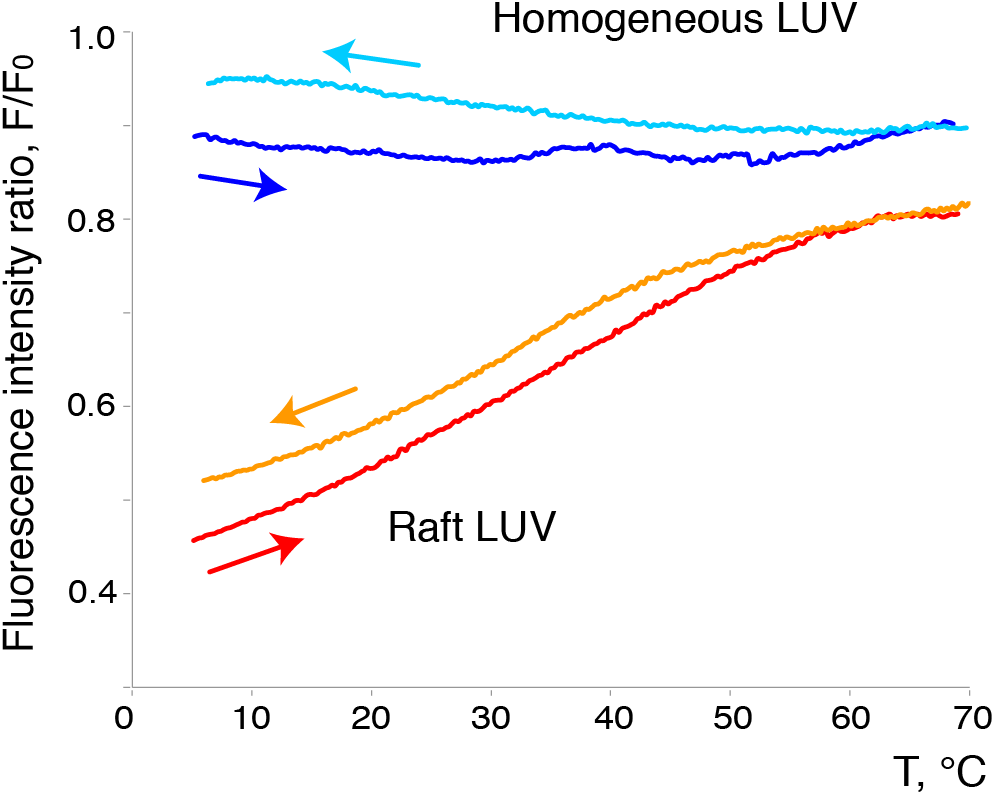
Non-raft localization of N-Ras C-terminal lipopeptide revealed by FRET to the disordered domain marker. Heating and cooling profiles of the homogeneous and raft LUV with Rhod-DOPE (acceptor; 2% mol) in the presence of the mant-labeled N-Ras C-terminal lipopeptide (donor; 0.1% mol). Fluorescence intensity ratio, F/F_o_, was calculated using mant emission of F and F_o_ samples, containing and lacking Rhod-DOPE, respectively. Each curve is an average of two independent samples. The F/F_o_ curves for homogeneous LUV showing high (yet relatively constant) values reflect difficulties with quantitative subtraction of light scattering caused by LUV in individual samples.

To confirm interpretation of rising F/F_o_ values as a signature of non-raft localization of the donor, we used another disordered lipid domain marker 1,2-dioleolyl-*sn*-glycero-3-phosphoethanolamine-N-5-dimethylamino-1-naphthalenesulfonyl (Dansyl-DOPE) and recorded the temperature dependence of F/F_o_ for dansyl group fluorescence (as a donor) in the presence and absence of the acceptor, Rhod-DOPE (Figure 4A). Dansyl-DOPE is localized in a disordered lipid phase along with Rhod-DOPE due to their unsaturated lipid chains (Figure 4B). A similar increasing trend in the temperature dependence of F/F_o_ values was observed confirming our conclusion of the localization of the mant-labeled C-terminal N-Ras lipopeptide outside of lipid rafts—readily accessible by acceptor fluorophores.

**Figure 4.**
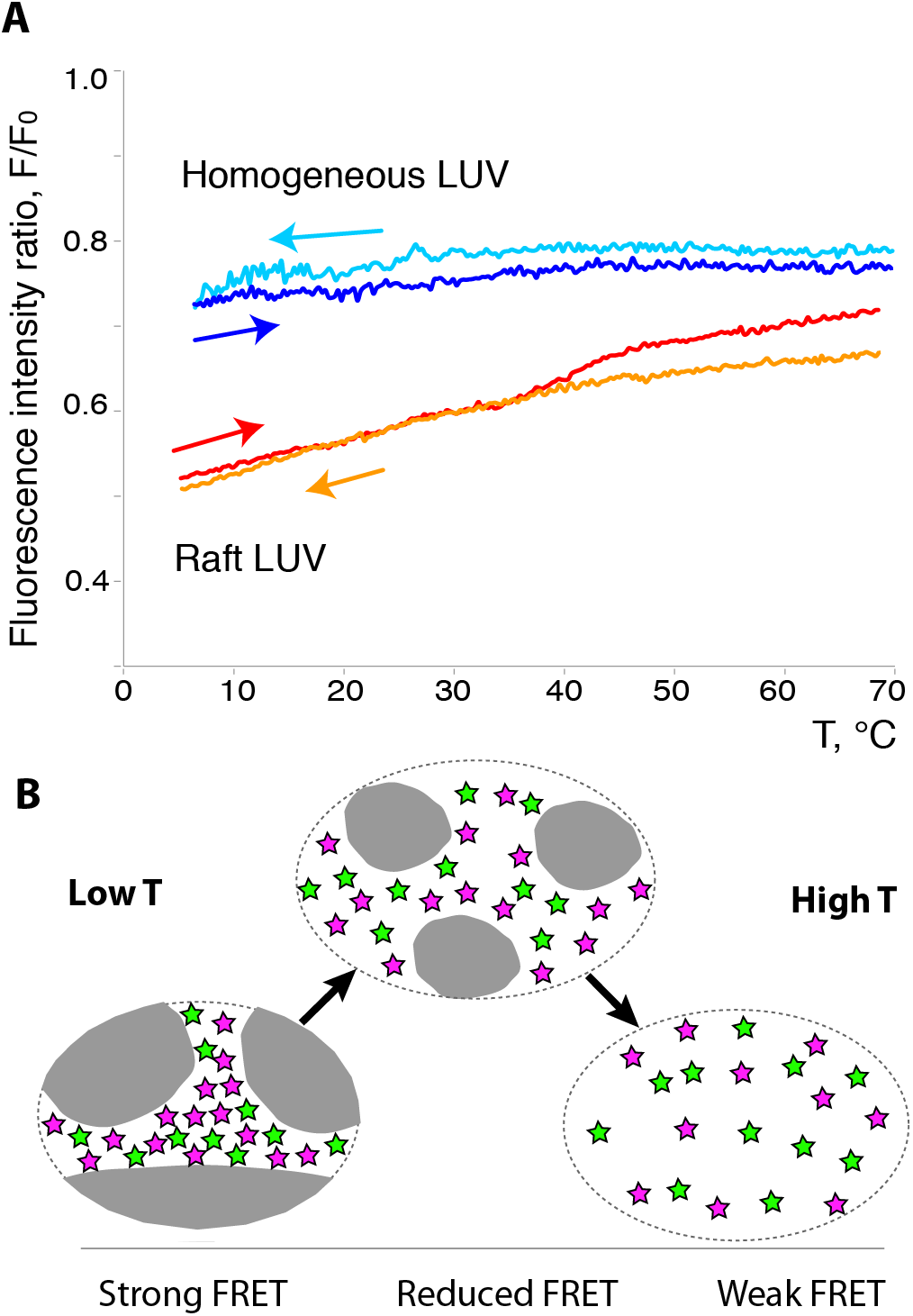
Disordered domain markers demonstrate increasing FRET upon reduction in raft size. (A) F/F_o_ temperature dependence for Dansyl-DOPE donor (0.1 % mol) incorporated into the homogeneous and raft-containing lipid bilayers containing Rhod-DOPE (2% mol). (B) Schematic drawing illustrating an increase in the average distance between donors (green stars) and acceptors (magenta stars) due to the melting of a raft phase (gray).

### 3. Test of the raft-boundary localization of the C-terminal N-Ras peptide

Experiments with mant-lipopeptide revealed that the lipopeptide is accessible to the acceptor fluorophore, Rhod-DOPE, at all times (Figure 3). However, these experiments could not discriminate the two possibilities: (1) lipopeptide uniformly distributed in the disordered lipid phase, and (2) lipopeptide concentrated at the boundary of the raft domains^*44, 60*^, because in both scenarios, the mant fluorophore will be easily accessible for quenching by rhodamine.

Localization of the lipopeptide at the raft boundary means the lipopeptide acts as a lineactant (attracted to the line between two-dimensional phases, analogous to surfactants populating surface separating three-dimensional phases) making the boundary more stable in its presence (i.e. line tension is reduced)^*61, 62*^. Stabilization of the raft boundary may be directly tested by evaluation of melting profiles in the presence of different concentrations of the lineactant. The lineactant facilitates increase of the total length of the boundary thus promoting breaking the existing rafts into smaller ones (destabilization of large rafts). Reduction in raft size will be detectable in FRET experiments with DPH and Rhod-DOPE, because DPH will be more effectively quenched by Rhod-DOPE in smaller rafts.

Figure 5 shows the heating profiles for raft LUV in the presence of increasing concentration of the lipopeptide (no fluorophore attached; see Supporting Information for estimates of the lipopeptide density at the raft boundary). Reduction in F/F_o_ values upon heating due to melting of lipid rafts occurs in a similar temperature range both in the absence and the presence of the lipidated N-Ras peptide. This observation implies that the raft boundary does not significantly attract the lipopeptide.

**Figure 5.**
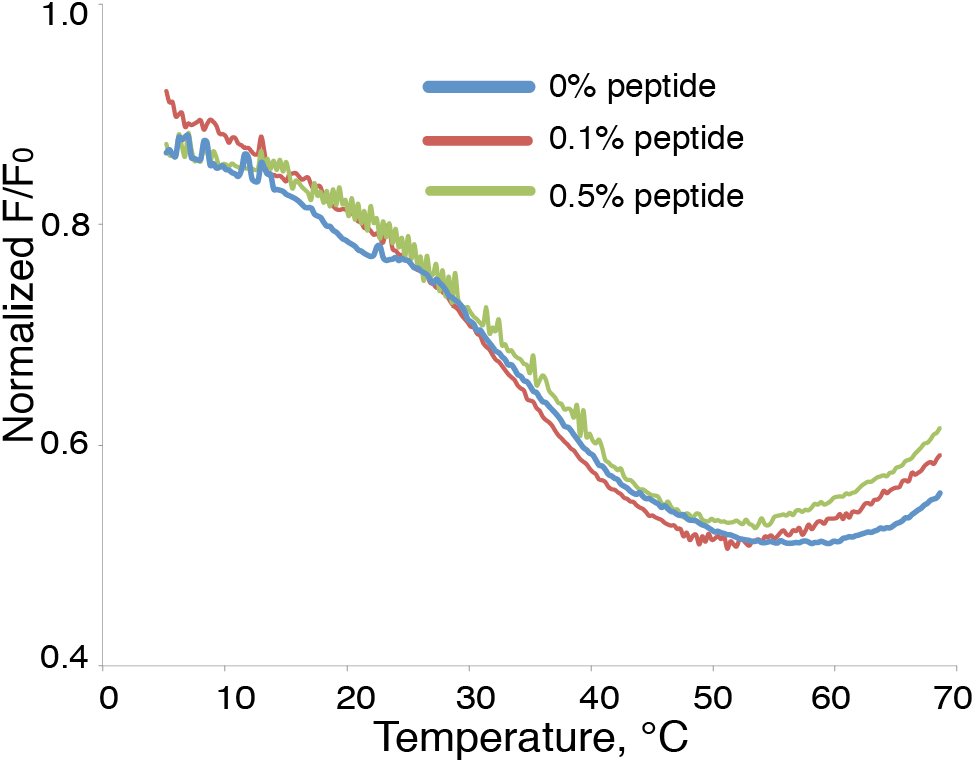
Test of a boundary localization of N-Ras C-terminal lipopeptide: heating profiles for the raft LUV with DPH and Rhod-DOPE and increasing concentration of non-fluorescent lipopeptide. The curves were shifted along Y axis to facilitate the comparison of the transition region.

### 4. Determination of the domain localization with time-domain fluorescence measurements

Analysis of the FRET donor distribution among ordered and disordered lipid domains in the heating/cooling experiments described above relied on a measurement of relative fluorescence intensities in the two samples with and without acceptor (F and F_o_), which required exactly matching concentrations of the donor. This is easy to accomplish for lipid mixtures that are made by taking accurate aliquots of fluorophore stocks, yet is very hard to achieve for the protein associated with LUV. The protein-LUV samples are made by mixing LUV with lipidated protein to allow for (always partial) spontaneous incorporation of lipoprotein. Protein density in a lipid bilayer is quite small and difficult to match between different samples. Determination of FRET through the *lifetime* measurements is superior to the intensity measurements in that it is *insensitive* to variations of donor concentration (with the downside: dramatically longer acquisition times). Lifetime measurements is a well-established method for characterization of environment of the fluorophore labels (for examples, see Bernsdorff et al. and Dong et al.^*63, 64*^) and lipid microdomains^*65–67*^.

Figure 6. A demonstrates this approach with the DPH-rhodamine FRET pair in homogeneous and raft-containing LUV. The homogeneous and raft mixtures lacking acceptor (F_o_ samples; green and black symbols) exhibited relatively invariable lifetimes throughout the full temperature range. In the presence of acceptor (F samples), the donor lifetime in a homogeneous sample (Figure 6.A, blue symbols) is reduced but remains relatively independent of the temperature. In contrast, the raft samples containing the acceptor (red symbols) reveal significant drop in DPH lifetime in a temperature range of the raft-melting transition. Figure 6.B demonstrates a corresponding increase in FRET efficiency reflecting greater quenching of DPH by Rhod-DOPE after rafts are melted (or reduced in the average size). This observation is in agreement with our results based on measurement of intensity in F and F_o_ samples in Figure 2 confirming presence of lipid raft nanodomains at low temperatures in the SM/POPC/cholesterol LUV samples.

**Figure 6.**
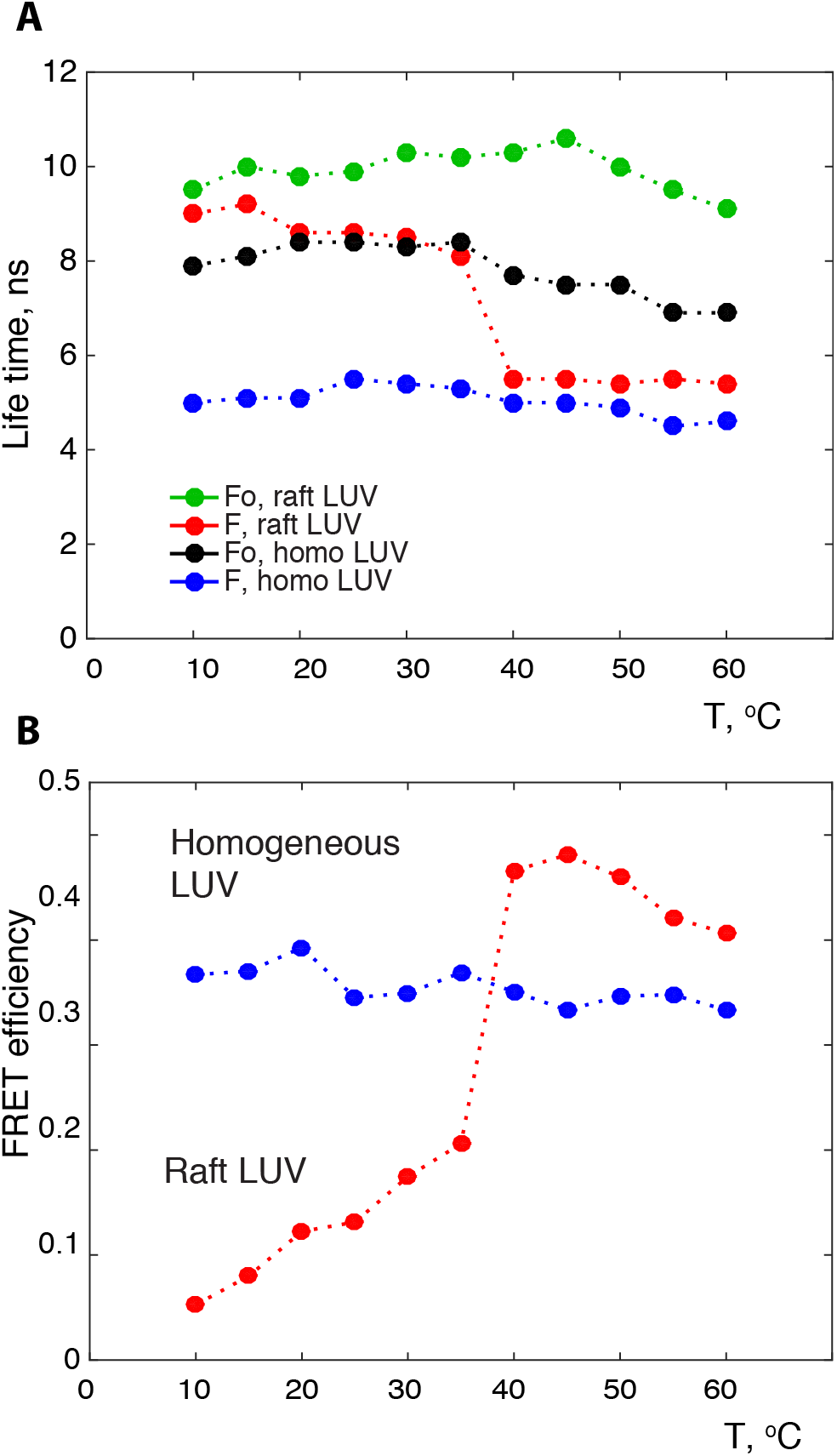
Raft stability in SM/POPC/cholesterol bilayers evaluated through time-domain fluorescence measurements. (A) Lifetimes of DPH fluorescence at different temperatures in homogeneous and raft-containing mixtures in the presence and the absence of acceptor Rhod-DOPE. (B) FRET efficiency calculated using Eq. 1 (see Materials and Methods) from lifetimes of DPH in panel A.

### 5. Preferential localization of N-Ras bound with fluorescent GDP and GTP-mimics

To determine domain localization of a N-Ras lipoprotein in raft membranes, we measured FRET between Ras-bound fluorescent derivatives of guanosine nucleotides as donors, and Rhod-DOPE localized in disordered lipid domains as an acceptor. To mimic a full-length N-Ras with the native posttranslational lipidation pattern (one palmitoyl and one farnesyl chain), we prepared a semisynthetic protein following protocols developed by Herbert Waldman group^*68–70*^. Protein samples were exchanged with mant-nucleotides and associated with LUV by overnight incubation. Lipoproteins tend to aggregate in aqueous buffers due to their hydrophobic lipid modifications; therefore, it was essential to ensure that any aggregated N-Ras that did not incorporate in LUV was removed before fluorescence measurements. Using size-exclusion chromatography, we achieved complete separation of N-Ras associated with LUV (eluted near exclusion limit of the column; >800 kDa) from the N-Ras aggregates (ca. 500 kDa) as well as from any unbound mant-nucleotides (Supporting Figure S1 and Table S2).

To establish the predominant localization of the N-Ras bound to mant-nucleotides, we determined efficiency of FRET between mant group and Rhod-DOPE in homogeneous and raft LUV. Since the thermal stability of Ras is relatively limited, we performed all experiments at low temperatures. Figure 7 shows FRET efficiencies calculated for mant in N-Ras-mGDP and N-Ras-mGppNp in homogeneous and raft LUVs (for a complete summary of the lifetime measurements see Table S1). Relatively high FRET efficiencies were observed for N-Ras-mGDP and N-Ras-mGppNHp associated with homogeneous lipid bilayers indicating significant energy transfer from mant fluorophore to rhodamine of Rhod-DOPE. This is an expected result as the donors are readily accessible to acceptors in the homogeneous bilayer (no domains). In raft LUV samples, N-Ras-mGDP exhibited very low FRET values indicating effective segregation of mant-labeled Ras-GDP from Rhod-DOPE at 5^°^C. These samples also exhibited relatively shorter life times, which might be explained by homotransfer^*71, 72*^ between mant groups due to protein clustering in the rafts^*22, 73, 74*^. In contrast, Ras-mGppNHp exhibited relatively high FRET values reporting on easy accessibility of mant to Rhod-DOPE.

**Figure 7.**
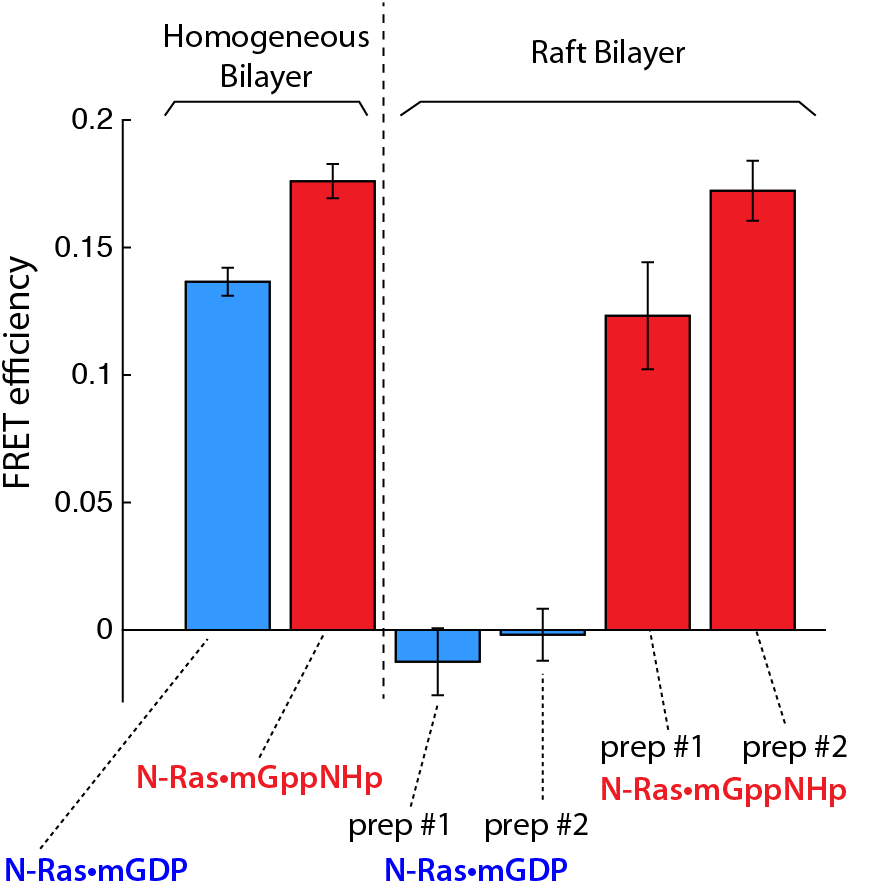
Efficiency of FRET between mant and Rhod-DOPE in samples of N-Ras-mGDP and N-Ras-mGppNHp at 5 ^°^C. Error bars indicate standard deviations from replicate lifetime measurements (for the numbers of replicates see Supporting Table 1). The raft LUV sample preparations were repeated to increase confidence in the result (indicated as prep #1 and #2, accordingly).

As an internal control, we attempted to convert raft samples into a “homogeneous” state by heating to 37^°^C when much of the raft phase is gone (see Figure 2). However, at this elevated temperature all homogeneous and raft LUV samples displayed near-zero FRET efficiencies suggesting that mant-nucleotides are completely separated from Rhod-DOPE. This separation might be due to dissociation of mant nucleotides from N-Ras upon heating considering long (1-4 hours) acquisition times of the TCSPC experiment and the weaker affinities of mant-nucleotides to Ras relatively to GDP and GTP. Therefore, we chose to limit our discussion in this paper to the low temperature at which N-Ras is most stable, and the rafts are relatively larger size^*50*^.

## Discussion

FRET analysis of N-Ras association with lipid nanodomains presented in Figure 7 revealed that N-Ras in the GDP-bound form (signaling-inactive state) concentrates in rafts but associates with the disordered phase when bound to a GTP-mimic. The raft localization of N-Ras-GDP is in agreement with *in vivo* observations of Matallanas et al., though the lipid domain preferences of N-Ras-GTP were not explored in their study^*42*^. It contrast, Roy et al. observed the G12V N-Ras clustered in cholesterol-dependent rafts (G12V mutation ensures that the proteins was predominantly bound to GTP) while the wild-type (GDP-bound) was found not clustered (remained outside of lipid rafts)^*43*^. Notable, that these research groups performed experiments on different types of cells: MDCK and COS-7^*42*^, and BHK^*43*^—variability in the lipid membrane composition and the presence of cell-type specific membrane proteins might be the cause for the observed opposite localization patterns.

In our study, we made it our goal to separate N-Ras interaction with lipids from possible interactions with other cellular membrane proteins. Choosing an adequate lipid raft model for an *in vitro* study is difficult because the cellular bilayers are asymmetric with their inner leaflet (where Ras proteins are) enriched with negatively charged phosphatidyl serine and completely lacking SM^*75, 76*^. The inner-leaflet mixture, however, cannot spontaneously form rafts, instead, raft formation must be triggered by the cross-leaflet interdigitation with the lipids of the outer side of plasma membrane (rich in SM)^*77, 78*^. Preparation of such asymmetric LUV was recently reported^*78, 79*^. However, performing experiments with asymmetric bilayers to study protein-nanodomain interactions is not straightforward as the lipid asymmetry is relatively short-lived—cells maintain it by a continuous action of lipid transporters, and the loss of lipid asymmetry is a signature of cell death^*80*^. LUV in our work and all other Ras reports were inherently symmetric that makes it difficult to evaluate as to which study made use of a more relevant lipid bilayer. Nicolini et al. used LUV made of DMPC/DSPC/cholesterol and did not recapitulate any of the *in vivo* observations^*44*^. Larsen et al. analyzed distribution of N-Ras C-terminal lipopeptide in DOPS/PSM/cholesterol bilayers and found it populating raft domains^*81*^. This is contrary to observation of Nicolini and others as well as our own data reported in this paper, which suggests that choice of the lipid system is crucial and far from settled.

In our study, the SM/POPC/cholesterol lipid mixture in equimolar ratio was used to create raft nanodomains most closely *mimicking the size* of cellular raft domains^*50*^, and helped reveal the “raft affinity switch” in the N-Ras macromolecule. Obtained data allow us to state that, while interactions with the cellular protein binding partners might be important for regulation of Ras domain preferences, the G domain itself controls the interaction with a raft phase, while C-terminal lipopeptide “pulls” the protein outside of the raft. We do not consider our evidence for nonboundary localization of the C-terminal lipopeptide particularly strong because we do not have a readily available positive control: a well-characterized lineactant that would serve as a calibration for the N-Ras peptide action in Figure 5. Therefore, our most accurate statement of N-Ras lipid domain preferences would be that G-domain in GDP-bound state is strongly attracted to the lipid rafts overcoming opposite preferences of the C-terminal lipopeptide; the Ras-raft interaction is weakened or absent in the GTP-bound form allowing the C-terminal lipopeptide to draw N-Ras outside of the raft (to the boundary or into the disordered membrane). Molecular dynamics simulation by Gorfe group determined that H-Ras C-terminal lipids favored localization at the raft boundary (due to palmitoyl chains favoring the ordered raft phase while the farnesyl lipid being excluded from it)^*82*^. Since N-Ras lipoprotein also has farnesyl and palmitoyl lipids, one might expect similar boundary localization for both the truncated C-terminal N-Ras lipopeptide and the full length N-Ras when bound to GTP.

Gorfe et al. identified a set of basic residues in H-Ras sequence that made specific contacts with lipid bilayer in molecular dynamics simulations—R128, R135, R169, and K170^*83*^. Abankwa and co-workers found these sites modulating Ras signaling function, and proposed that conformations of these residues and overall orientation of the G domain must be affected by the nucleotide-binding site via an allosteric coupling mechanism^*51, 52,84*^. These positively charged sites are presented in N-Ras by conservative substitutions: K128, K135, K169, while K170 is identical. At about the same time, using spin-relaxation NMR measurements we reported that G domains of H-Ras and K-Ras possessed global conformational exchange dynamics connecting the effector interface of Ras with the rest of the molecule^*85, 86*^. We demonstrated that the novel ion-binding pocket on the membrane-facing side of the G domain in H-Ras described by Buhrman and coworkers^*87*^ was thermodynamically coupled to nucleotide binding site while being at nearly 20 Å distance away: its affinity for a divalent ion changes by a factor of five upon replacement of GDP with the GTP-mimic^*88*^. Due to high sequence identity of the G domains we reasonably expect the N-Ras to possess the same property.

Yet, the plausible molecular mechanism of the *lipid-domain recognition* by Ras isoforms has not been resolved experimentally. Instead, Werkmuller et al. provided evidence that G domains of N and K-Ras are experiencing similar rotational freedom next to the membrane regardless of the bound nucleotide or a type of the lipid bilayer (raft or homogeneous)^*89*^. Similarity of rotational diffusion in all conditions implied that their G domains do not interact with the lipid other than through a C-terminal lipidated peptide acting as a tether (however, it was not clear if the Ras-LUV samples were separated from aggregated lipidated Ras or the authors simply assumed 100% binding and zero contribution from protein aggregates). This is why a view that other cellular proteins like galectins interact with the GTP-bound Ras and cause redistribution between lipid domains remains an attractive alternative^*38*^. Yet, N-Ras was not found to bind galectins and, most importantly, galectin 1 (interacting with H-Ras) is a cytosolic protein^*90*^ *recruited by* H-Ras to the plasma membrane^*41*^ making it less likely to be the domain recognition driver. In our experiments, the N-Ras∂mGppNHp demonstrated association with rhodamine-labeled disordered domains while N-Ras∂mGDP concentrated in rafts—all in the absence of other cellular components, which forces us to reconsider the role of a G domain in the lipid raft recognition by Ras proteins.

Finally, we need to make an important cautionary note on the differences that may be reported in studies using bulk fluorescence measurements of LUV vs. confocal microscopy experiments (including single-molecule tracking). The confocal microscopy, by design, involves focusing of a very intense laser light into a very small area to ensure effective excitation. This mode of observation was demonstrated to create artifacts due to overly intense illumination, particularly, when observing Ras, which is a tyrosine-rich molecule prone to irreversible photoactivated cross-linking^*91*^. Earlier, we established that the light intensity used in a solution fluorescence measurement in a conventional spectrofluorometer did not lead to cross-linking artifacts in a highly homologous Ras construct^*92*^—therefore, oxidative cross-linking of N-Ras was not expected to negatively affect the results of the current report.

## Conclusions

In this study, we demonstrated that N-Ras lipoprotein changes its lipid nanodomain preferences in a nucleotide-dependent manner in the absence of other membrane proteins (in a model lipid membrane). *The signaling-inactive, GDP-bound N-Ras was found to have preferential affinity for lipid rafts. N-Ras in its activated conformation (bound to GTP-mimic) was localized at the raft boundary or in a disordered lipid phase*. Thus, we established that the specific nanodomain preference is an intrinsic property of the full-length N-Ras lipoprotein, which may further be modulated by specific protein-protein interactions in the cell.

## Materials and Methods

Materials and Methods section in its entirety is included in Supporting Information.

## Acknowledgment

This work was funded by NSF (CHE-1508677) and NIH (R01-HL112639-04) to RR; the Regular Research Grant 2012 from Committee on Research (COR), Marquette University, to ELK. AS acknowledges support from Arthur J. Schmitt Leadership Fellowship 2016-2017; ARG acknowledges Eisch Research Fellowship during the academic year 2016-2017. ELK acknowledges Summer Faculty Fellowship 2012 from Committee on Research (COR), Marquette University. Authors are grateful to Dr. Erwin London and Dr. Alemayehu Gorfe for critical reading of the manuscript and helpful discussions.

## Supporting Information

Complete Materials and Methods section; Supporting Tables 1 and 2 with lifetime and size-exclusion data; Supporting Figures; estimates of protein surface densities.

---

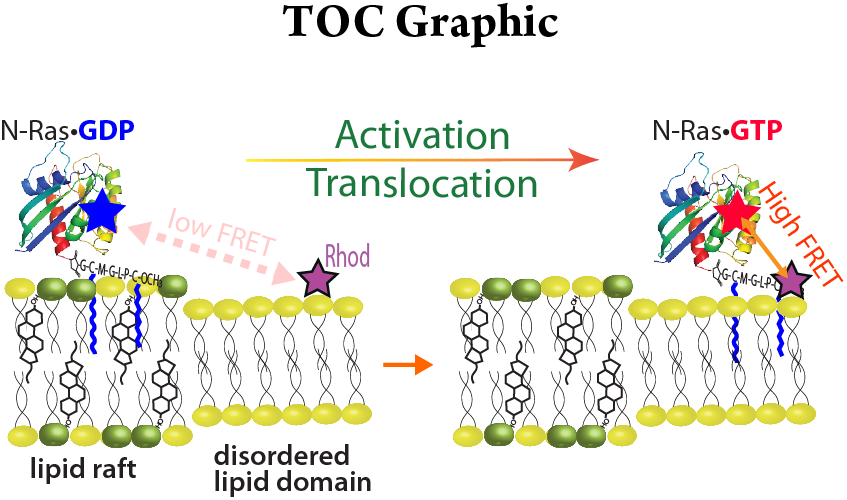

## Supporting Information

### Materials and Methods

#### Materials

All chemicals and solvents were obtained from commercial suppliers (Thermo Fisher and VWR) and used without further purification. Fmoc-protected amino acids, 2-(1H-benzotriazol-1-yl)-1,1,3,3-tetramethyluronium hexafluorophosphate (HBTU) and hydroxybenzotriazole (HOBt) were obtained from Advanced Chem Tech. 1,6-Diphenyl-1,3,5-hexatriene (DPH), cholesterol, Triton X-114, and trans, trans-farnesyl bromide and trimethylacetic anhydride were obtained from Sigma Aldrich. 6-Maleimidohexanoic acid was obtained from Alfa Aesar. *N*-Methylanthranilic acid was obtained inform TCI America. Chicken egg sphingomyelin (SM), 1-palmitoyl-2-oleoyl-phosphatidylcholine (POPC), 1,2-dioleoylphosphoethanolamine-N-(lissamine rhodamine B sulfonyl) (Rhod-DOPE), 1,2-dioleoyl-sn-glycero-3-phospho-(1’-rac-glycerol) (sodium salt) (DOPG), 1,2-dioleoyl-sn-glycero-3-phosphoethanolamine-N-(5-dimethylamino-1-naphthalenesulfonyl) (ammonium salt) (Dansyl-DOPE) were purchased from Avanti Polar Lipids (Alabaster, AL). (2’-(or-3’)-O-(N-Methylanthraniloyl) guanosine 5’-diphosphate (mant-GDP) was obtained from Biolog Lifescience Institute 2’-/3’-O-(N’-Methylanthraniloyl) guanosine-5’-O- [(β, γ-imidotriphosphate] (mant-GppNHp) was obtained from Life Technologies.

#### Protein constructs

The full-length gene of the wild-type *N*-Ras was a gift of Dr. Robert Deschenes, University of South Florida. For bacterial expression, N-Ras gene was subcloned into the pET vector (EMD Millipore, Billerica, MA). The truncated N-Ras construct ending with the cysteine 181 was prepared by introducing a stop codon in place of the methionine 182 codon. The C118S mutation was introduced to avoid possible side reactions between the only exposed cysteine on the G domain and maleimido group of the lipidated peptide^*93*^. All mutagenesis steps were performed using the QuikChange Site-Directed Mutagenesis Kit (Life Technologies, Grand Island, NY).

#### Protein preparation

Expression and isolation NRas-C118S-181 was performed as described earlier for a similar construct of H-Ras with little modifications^**92*, 94*^. Final yield was approximately 0.5-2 mg of 95% pure protein from each liter of the expression medium.

#### Preparation of the lipidated peptides

Fmoc-protected farnesylated cysteine was prepared as described^*95*^. For the preparation of Fmoc-protected hexadecylated cysteine, cysteine was alkylated in the presence of 1,1,3,3-tetramethylguanidine followed by introduction of Fmoc-group using Fmoc *N*-hydroxysuccinimide ester in the presence of triethylamine^*96*^.

All lipidated peptides were prepared in a manual solid-phase reactor following published Fmoc chemistry protocols^*70, 97*^. In brief, 200-350 mg of 4-Fmoc-hydrazinobenzoyl AM Nova Gel (Novabiochem) (substitution 0.64 mmol/g) were used to couple 4 eq Fmoc-protected farnesylated cysteine in the presence of 4 eq HBTU and 4 eq of HOBt. Fmoc-group was removed by treatment with 20% pyridine in DMF. Each subsequent amino acid was coupled in a similar manner. In case of maleimide- or mant-containing lipopeptides, the 6-maleimidohexanoic acid or *N*-methylanthranilic acid were introduced at the last step of coupling, respectively. To increase the purity of the final lipopeptide, Double coupling followed by capping with 10% of trimethylacetic anhydride in DMF was used for introduction of non-lipidated amino acids. Finally, the peptides were cleaved off the resin by treatment with 0.5 eq Cu(OAc)_2_, 30 eq pyridine, 15 eq acetic acid and 250 eq methanol. We used hexadecyl group in place of palmitoyl for improved stability of the modification^*70, 97*^. The synthesized peptides were purified using the RP-HPLC-C4 column (Phenomenex). The molecular mass of the pure product was confirmed using matrix-assisted laser desorption ionization (MALDI) mass and tandem liquid chromatography/mass spectrometry (LC-MS) (See Supporting Information for yields and LC-MS data).

#### Conjugation reaction and purification of lipidated Ras construct

Conjugation reaction was performed in Triton X-114 solution. This non-ionic detergent undergoes phase separation in aqueous solutions at temperatures above 30°C. Hydrophobic or lipid-modified proteins stay in detergent phase upon separation. The Triton X-114 solution was prepared prior to the reaction to achieve the final concentration of about 30 g/L as described^*98*^. Purified N-Ras-C118S-181 protein (10 mg/ml) was dialyzed against 20 mM Tris, pH 7.4, 5 mM MgCl_2_ (coupling buffer) to remove DTT and excess NaCl, which may interfere with the coupling reaction. Lipidated peptide (1 mg) was solubilized in 50 μL of methanol. The 1 ml of Triton X-114 solution was added to the methanolic solution and sonicated in the ultrasonic bath until the peptide was fully dissolved. Then, 1ml of the obtained peptide solution was added to 1 ml of the dialyzed protein sample (1.5:1 mole peptide/mole Ras). The reaction mixture was flushed with argon and incubated overnight at 4°C on a rotator. Next day the mixture was spun at 4^*o*^C for 12 min at 10,000 g and the supernatant was transferred to a clear plastic tube. The reaction was quenched by addition of equal volume of 1 mM DTT in cold 20 mM Tris, pH 7.4, 5 mM MgCl_2_. To allow for separation of an unreacted protein from the semisynthetic product, the mixture was heated to 37°C until solution became cloudy due to separation of the detergent-rich phase and centrifuged for 10 min at 3500 g with the centrifuge brakes turned off to avoid vibration. The detergent-rich and aqueous phases were separated by aspiration (the aqueous phase was on the top). The detergent phase was combined with a fresh DTT-containing coupling buffer; the aqueous phase was combined with Triton X-114 solution, respectively. The procedure was repeated 3 times. All detergent phases were combined and diluted 10-fold with the cold coupling buffer with 1 mM DTT. To selectively remove the detergent we used Amberlite XAD-2 (Supelco) beads as an absorbent. The 3 g (dry weight) of beads were soaked in methanol followed by a thorough wash with a coupling buffer. The wet beads were added to the 7 ml of diluted detergent phase to provide 20-fold detergent-binding capacity relatively to the amount of Triton X-114 (considering stated 3 g of beads per 0.1 g of Triton) and incubated overnight at 4°C on a rotator. The mixture was filtered and assessed by absorption at 254 nm to verify that concentration of Triton X-114 was reduced below 0.1 mM. Aqueous phases were pooled together to evaluate fraction of Ras protein that did not participate in the conjugation reaction.

To confirm the success of the conjugation reaction, we performed SDS-PAGE and MALDI-TOF analysis of the protein samples before and after conjugation. Figures S5 and S6 show an expected increase in mass of 1315 Da indicating that Ras was successfully conjugates to the lipidated peptide.

#### LUVs preparation

Lipids and their fluorescent derivatives were dissolved in chloroform (with the exception of DPH, which was dissolved in ethanol) and stored at -20°C. The concentrations of fluorescent lipids were determined by absorbance using ε(Rhod-DOPE) = 88,000 M^−1^cm^−1^ at 560 nm, ε(DPH) = 84,800 M^−1^cm^−1^ at 352 nm, and ε(mant-GDP) = 22,600 M^−1^cm^−1^ at 360 nm.

Lipid unilamellar vesicles, LUV, were prepared by extrusion following published protocols^*50, 99*^. Lipids, fluorophores and peptides were mixed in glass vials. Rhod-DOPE was added to 2% (mol of total lipid) to create F samples. To make the F_o_ samples, Rhod-DOPE was substituted by 2% DOPG to remove acceptor fluorophore but maintain the negative charge of the bilayer. Donor fluorophores DPH, dansyl-DOPE, and mant-lipopeptide, were added to F and F_o_ samples to 0.1% mol of total lipid, respectively. The chloroform was evaporated under a stream of nitrogen gas, and the dried lipid film was rehydrated in 20 mM Tris, pH 7.4, 5 mM MgCl_2_, 150mM NaCl buffer at 70°C. The LUV were formed by extrusion: the solution with a total lipid concentration of 0.5 to 1 mM was passed 21 times through a polycarbonate filter with a pore size of 0.2 μm (Avanti Polar Lipids). The temperature of the exturder block was maintained at 75 ^*o*^C. LUV lacking fluorescent probes was also prepared for subtraction of background fluorescence.

#### Preparation of Ras-LUV samples

Fluorescent Ras-GDP and Ras-GTP complexes were prepared using the (2’-(or-3’)-O-(N-methylanthraniloyl) guanosine 5’-diphosphate, mGDP, and the slowly hydrolysable GTP mimic 2’/3’-O-(N-methyl-anthraniloyl)-guanosine-5’-[(β,γ)-imido] triphosphate, mGppNHp, respectively. To prepare N-Ras-mGDP or N-Ras-mGppNHp associated with LUV, the lipidated Ras samples were subject to the nucleotide exchange followed by association with LUV and chromatographic separation as described in the following.

For introduction of mGDP, the GDP nucleotide associated with the GTPase site in Ras was displaced with the mGDP by mass action using the EDTA-assisted method^*100*^. In brief, the magnesium ions (5 mM MgCl_2_) in the 40 *μ*M protein samples were chelated with 6 mM EDTA; the 10 mM DTT (final concentration) was added to protect the protein cysteines from oxidation. Nucleotide exchange was started by addition of mGDP to 0.8 mM. The reaction mixtures were incubated for 2 hours at room temperature. Exchange reaction was quenched by addition of 10 mM MgCl_2_. Association with LUV was initiated by mixing 180 *μ*L of the quenched reaction mix with 400 *μ*L of LUV solution (1 mM total lipid). The mixtures were incubated overnight at 4°C and separated by passing through Superose 6 Increase 10/300 GL column (GE Healthcare) equilibrated with 30 mM Tris, pH 7.4, 5 mM MgCl_2_, 150 mM NaCl, 1 mM DTT buffer. Size-exclusion elution profiles were monitored by tyrosine and mant fluorescence for protein, and rhodamine fluorescence for LUV. Fractions corresponding to the hydrodynamic radius of LUV (near exclusion limit of the column) were used directly in fluorescence measurements. To verify that the LUVs do not capture or bind free mant-nucleotides, the “mock” exchange reaction was performed with the protein omitted from the mix. Subsequent separation by size-exclusion demonstrated that all mant-nucleotides elute in the full volume of the Superose 6 column corresponding to their small molecular weight. In the separate size-exclusion experiments, we determined that the non-lipidated Ras does not bind to LUV, while the aggregated lipidated Ras (by itself; no LUV added) elutes outside of the LUV size range allowing for effective separation of unbound lipidated Ras from Ras-LUV conjugates (Supporting Figure S1). Elution volumes of all samples are given in Supporting Table 2.

Nucleotide exchange for mGppNHp was performed through mass action facilitated by enzymatic cleavage of displaced GDP^*101*^. In short, the 16 *μ*M protein solution was dialyzed against nucleotide-exchange buffer containing 50 mM TRIS, pH 8.0, 5 mM DTT, and 200 mM (NH_4_)_2_SO_4_. Nucleotide exchange was started by addition of mGppNHp to 1 mL of protein solution to reach 2x molar excess over the protein. Shrimp alkaline phosphatase (Fermentas) was added (10 units) to hydrolyze released GDP molecules. Exchange was allowed to proceed for 2 hours at room temperature. Obtained Ras-mGppNHp sample was dialyzed against 20 mM TRIS, pH 7.4, 1 mM DTT, 5 mM MgCl_2_ buffer, associated with LUV, and separated from unbound lipidated Ras by size-exclusion chromatography as described for Ras-mGDP above.

#### Confocal microscopy of supported lipid bilayers

Supported lipid bilayers were created using raft LUV and observed with Nikon Perfect Focus Ti-E inverted research microscope using standard laser and filter sets. NBD-DPPE (1,2-dipalmitoyl-sn-glycero-3-phosphoethanolamine-N-(7-nitro-2-1,3-benzoxadiazol-4-yl)) was used as a lipid raft marker, Rhod-DOPE (1,2-dioleoyl-sn-glycero-3-phosphoethanolamine conjugated with lissamine rhodamine B) was used as disordered domain marker at 0.5% (molar) each^*10*^. To create supported lipid bilayers, LUV solutions were applied to clean glass slides and carefully rinsed to remove unbound LUV while keeping glass surface covered with solution at all times^*102*^. NBD fluorescence image (in green) was overlayed on rhodamine fluorescence image (in red) to create Figure 1.

#### Fluorescence spectroscopy

Measurements of steady-state and time-resolved fluorescence were performed using the Photon Technology International QM40 QuantaMaster system equipped with Pico-Master 1 time-correlated single-photon counting unit (HORIBA Scientific, Edison, NJ). A four-position Peltier-based Turret 400 (Quantum Northwest, Shoreline, WA) allowed for simultaneous temperature control and observation of up to four replicates for each sample condition. DPH, mant, and dansyl fluorophores were excited at 350, 360, and 340 nm, and their fluorescence was detected at 425, 440, and 520 nm, respectively. In heating and cooling experiments, temperature change rate was set 0.5°C. Temperature differences between the cells were directly tested with a digital thermometer and did not exceed 0.5°C. Time-domain fluorescence measurements of DPH and mant were done using the 365 nm LED with a pulse width of approx. 1 ns. The slit widths of the emission monochromator were adjusted in a range from 1 to 8 nm to maintain the TCSPC counting rate below 2%. The instrument response function, IRF, was recorded using a solution of a generic scatterer. Time-domain fluorescence decays were analyzed using *DecayFit* software (kindly shared by Søren Preus; available from www.fluortools.com). The three-exponential decay function was used to model the decay of the fluorophore fluorescence as well as a contribution of excitation light scattered by LUV in the samples. The *DecayFit* software subtracted scattered light using variable contribution of the IRF while the rest of scattered photons were accounted for by the fast-decaying component of the three-exponential model (yielding a sub-nanosecond life time). The second component of the model with the life time on the order of 5-8 ns was used as representative of the fluorophore life time in the samples. The third life time, typically—on the order of 20-40 ns, contributed very small percentage of signal (<5%) and, therefore, was ignored in the FRET analysis. The life time constants of the second component of the model obtained by least-squared fits to the data are summarized in Supporting Table 1 and referred to in this work as the “life time constants” of the samples. FRET efficiency was calculated using Eq. 1:

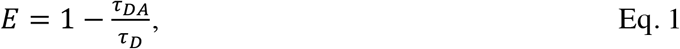

where *τ_DA_* and *τ_D_* are the life times of donor in the presence and the absence of the acceptor (F and F_o_ samples), respectively. Standard deviation of FRET efficiency was estimated assuming independent errors of life times^*103*^ expressed by the Eq.2:

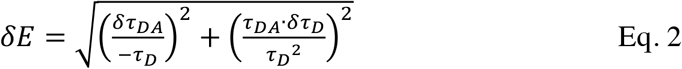

### Supporting Tables and Figures

**Table S1.** Lifetimes of mant fluorophore in N-Ras lipoprotein samples loaded with mGDP and mGppNHp associated with homogeneous and lipid raft LUV. Replicates were acquired by measurements of aliquots of the same sample in four separate positions of the sample turret of the spectrofluorometer (recorded one after another). Standard deviations and number of replicates are given in parentheses. FRET efficiency values and their standard deviation were calculated using Eq. 1 and Eq. 2. Raft LUV sample preparations were repeated starting from fresh mant-nucleotide exchange reaction followed by coupling to freshly extruded LUV and by gel filtration to ensure reliability of the measurement results (shown as preparations 1 and 2).

| N-Ras•mGDP |  |  |  |  |
| --- | --- | --- | --- | --- |
| Lipid mix | Homogeneous LUVs | Raft LUVs |  |  |
| Temperature | 5 °C | 5 °C<br>preparation 1 | 5 °C<br>preparation 2 | 16 °C<br>preparation 2 |
| Lifetime $F_0$, ns | 7.47<br>(0.01, 4) | 4.50<br>(0.04, 4) | 5.28<br>(0.05, 4) | 5.16<br>(0.13, 2) |
| Lifetime $F$, ns | 6.45<br>(0.04, 3) | 4.56<br>(0.05, 4) | 5.29<br>(0.02, 4) | 5.17<br>(0.15, 2) |
| FRET Efficiency | 0.137<br>(0.005) | -0.012<br>(0.013) | -0.002<br>(0.010) | -0.002<br>(0.04) |
| N-Ras•mGppNHp |  |  |  |  |
| Lipid mix | Homogeneous LUVs | Raft LUVs |  |  |
| Temperature | 5 °C | 5 °C<br>preparation 1 | 5 °C<br>preparation 2 |  |
| Lifetime $F_0$, ns<br>($\tau_D$) | 8.92<br>(0.04, 4) | 7.87<br>(0.15, 4) | 6.56<br>(0.04, 2) | |
| Lifetime $F$, ns<br>($\tau_{DA}$) | 7.35<br>(0.05, 3) | 6.90<br>(0.1, 4) | 5.43<br>(0.07, 2) | |
| FRET Efficiency | 0.176<br>(0.007) | 0.123<br>(0.021) | 0.172<br>(0.012) |  |

**Table S2.** Elution volumes and hydrodynamically equivalent size (spherical protein standard) for gel-filtration of Ras-LUV samples on Superose 6 Increase 10/300 GL column (GE Healthcare).

| **Sample** | **Elution volume** | **Corresponding size of a globular standard** |
| --- | --- | --- |
| N-RasC118S-181 (no lipid) | 25 ml | <30 kDa |
| Homogeneous LUV | 11 ml | >800 kDa |
| raft LUV | 11 ml | >800 kDa |
| N-Ras-mGDP, lipidated, no LUV added | 15-16 ml | ~500 kDa |
| LUV with N-Ras-mGDP | 11 ml | >800 kDa |
| LUV with N-Ras-mGppNHp | 11 ml | >800 kDa |

**Figure S1.**
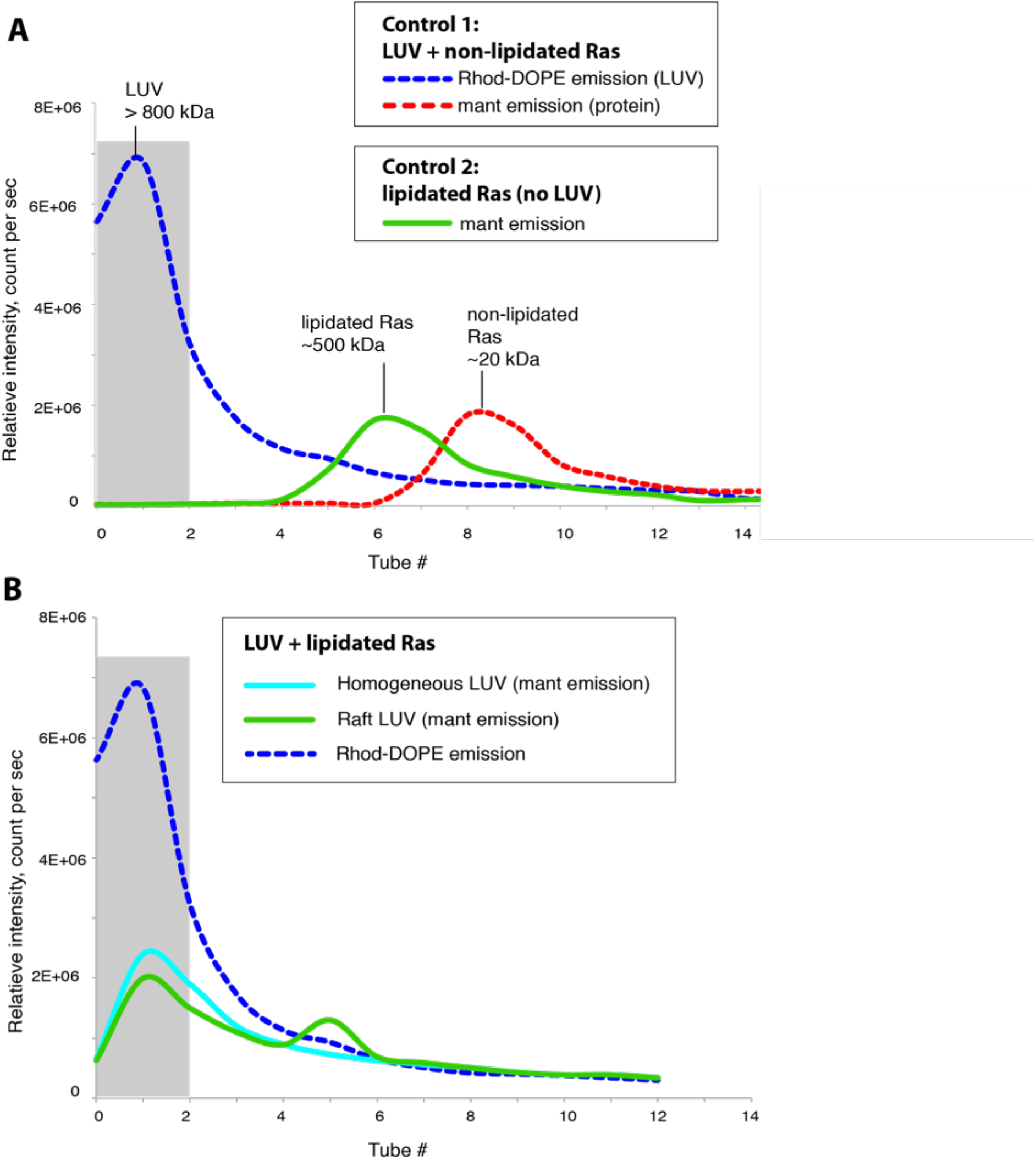
Representative elution profiles of control samples (**A**) and Ras-LUV conjugate samples used in FRET measurements (**B**). The LUV elution was followed using rhodamine emission at 590 nm (excited at 560 nm), the protein elution—using mant-nucleotide emission at 440 nm (excited at 360 nm). In **A**, Control 1 sample contained a mixture of LUV and non-lipidated Ras; Control 2 had lipidated Ras but no LUV. In **B**, a rhodamine emission profile was identical for both Ras-LUV conjugation reactions—shown by a blue dashed line. Shaded area represents the collection range for the Ras-LUV conjugate sample.

**Figure S2.**
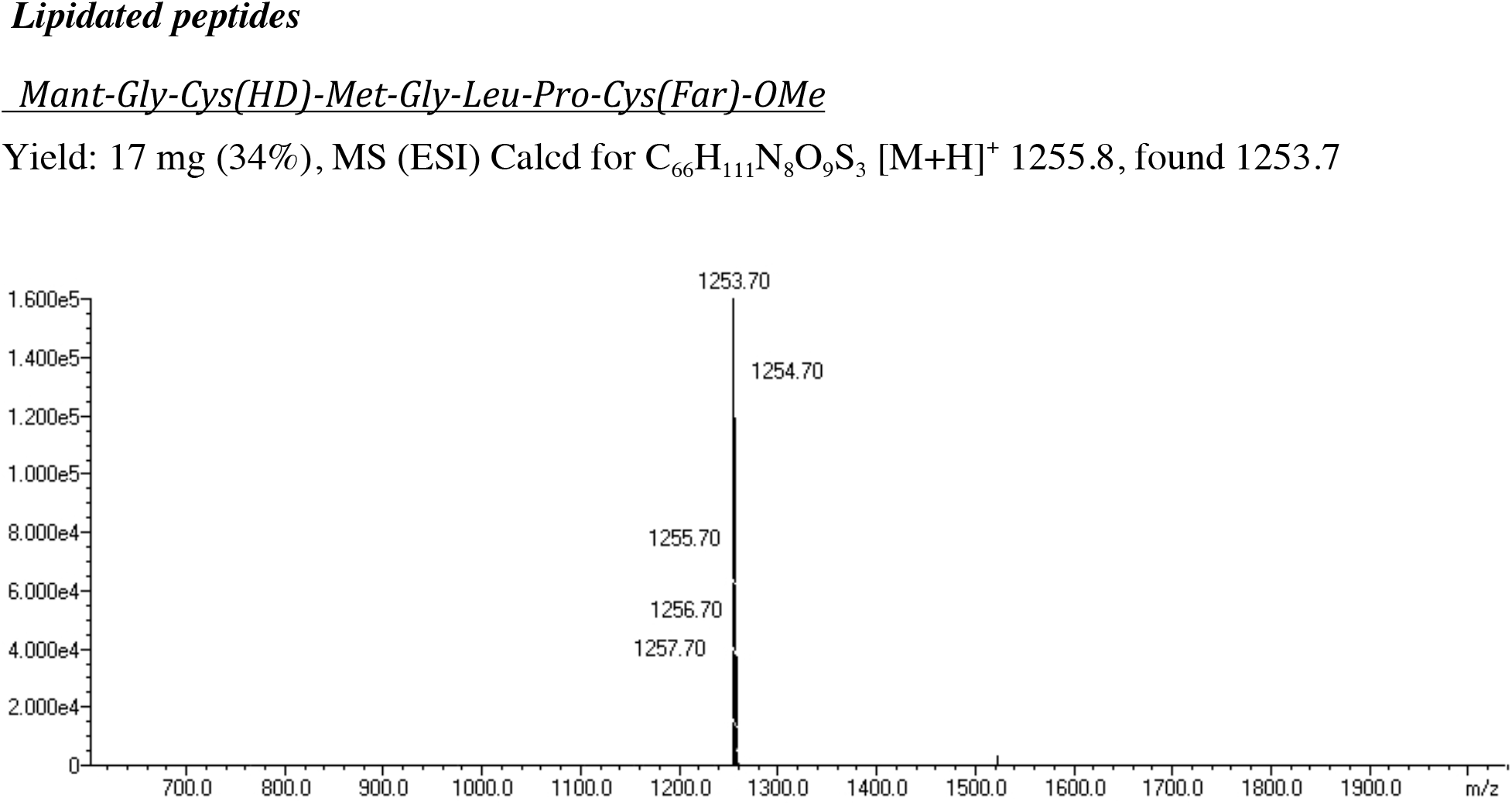
Mass spectrum of unlabeled lipopetide

**Figure S3.**
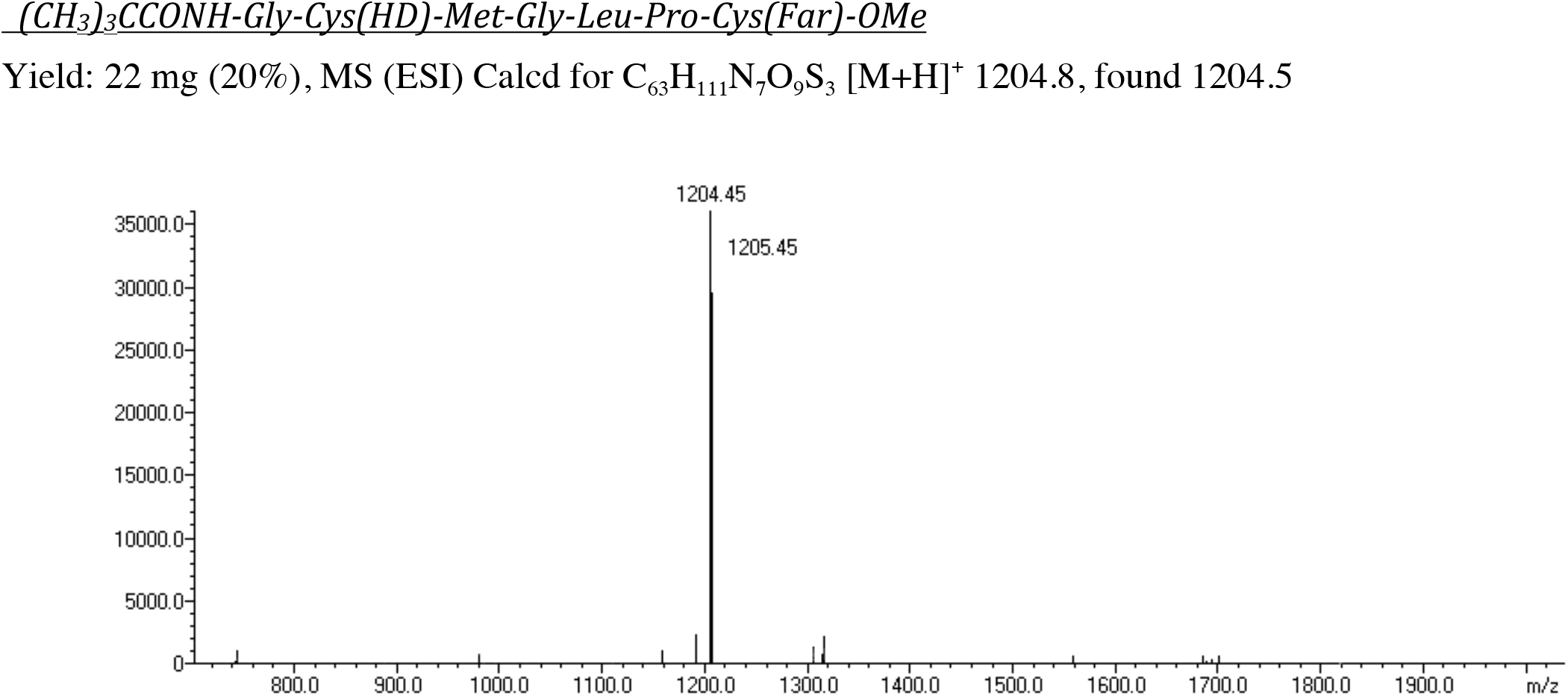
Mass spectrum of mant-labeled lipopetide

**Figure S4.**
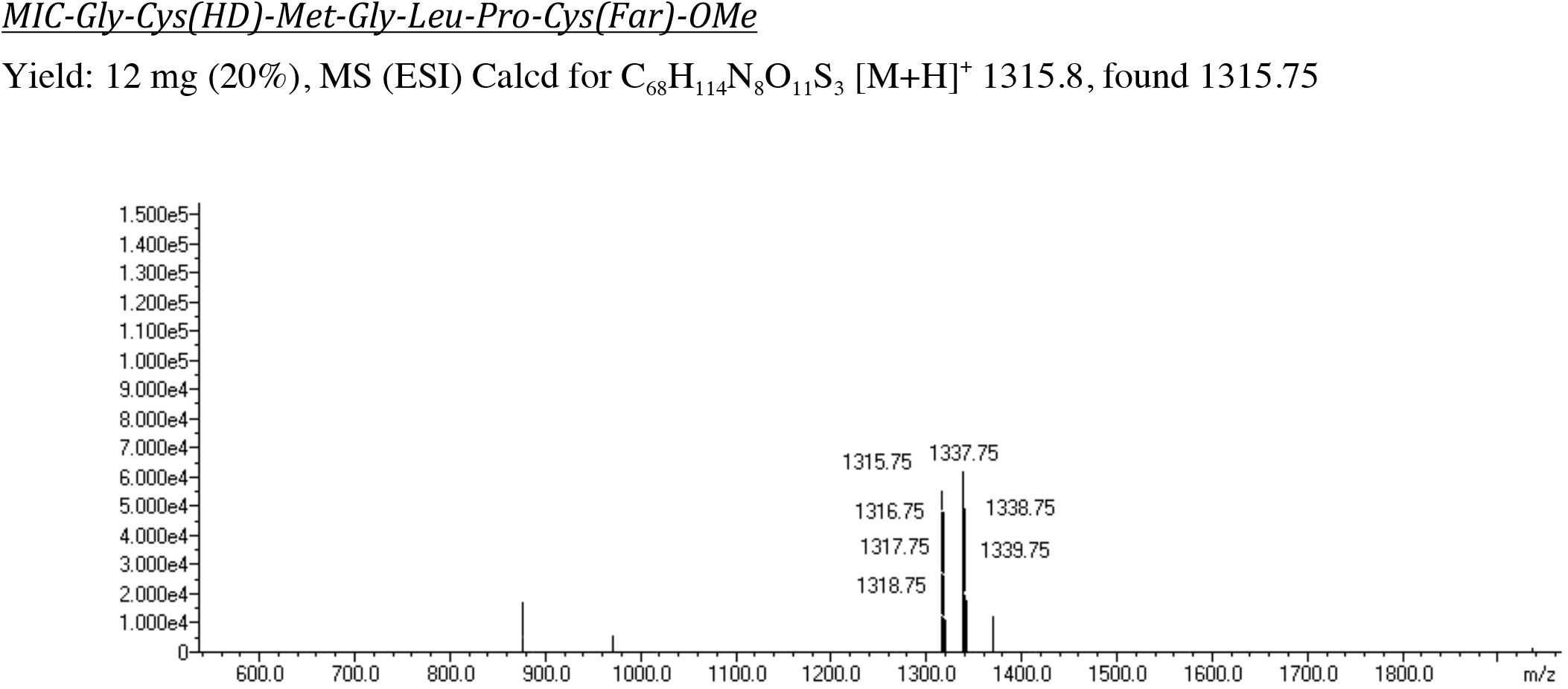
Mass spectrum of MIC-lipopetide for conjugation with N-Ras

**Figure S5.**
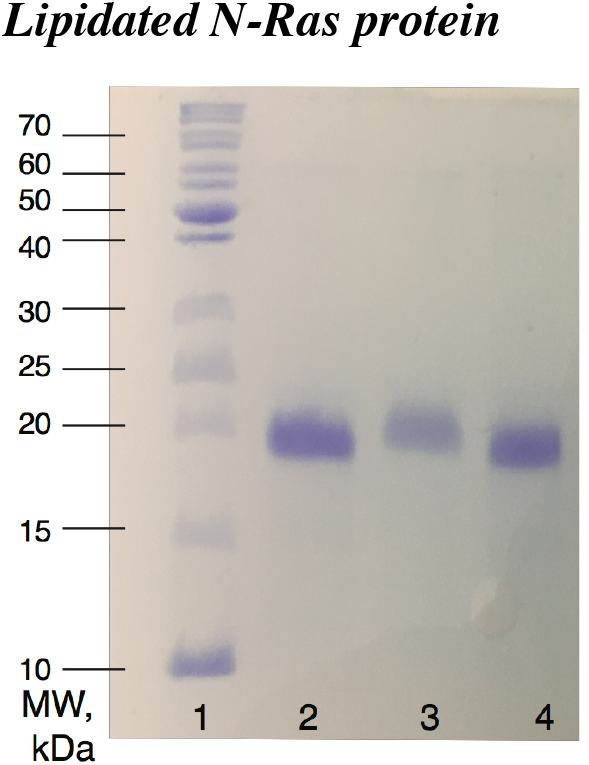
SDS-PAGE of the C118S N-Ras protein before (lanes 2 and 4) and after conjugation with lipidated peptide (lane 3).

**Figure S6.**
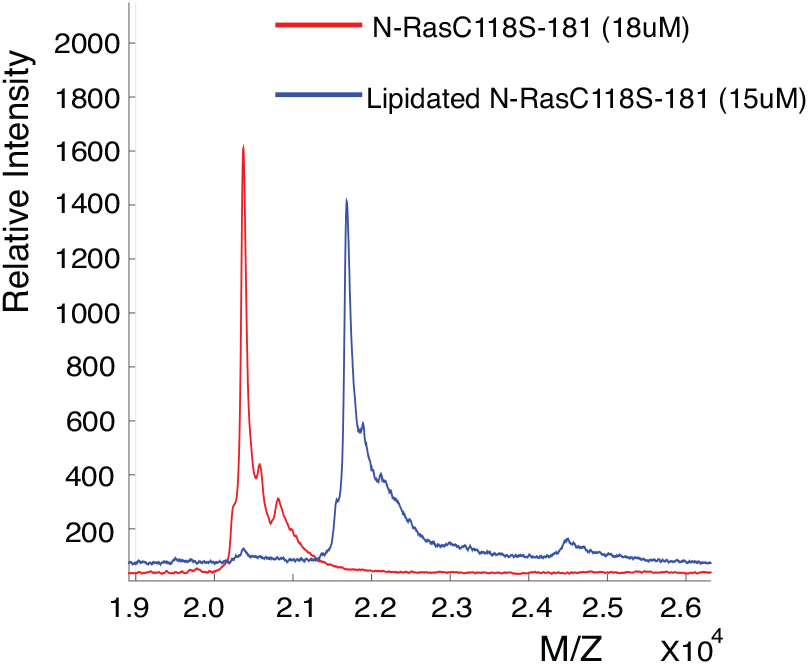
MALDI-TOF spectra of C118S N-Ras protein before and after conjugation with lipidated peptide. The difference in masses of the major peaks is 1315 Da.

**Figure S7.**
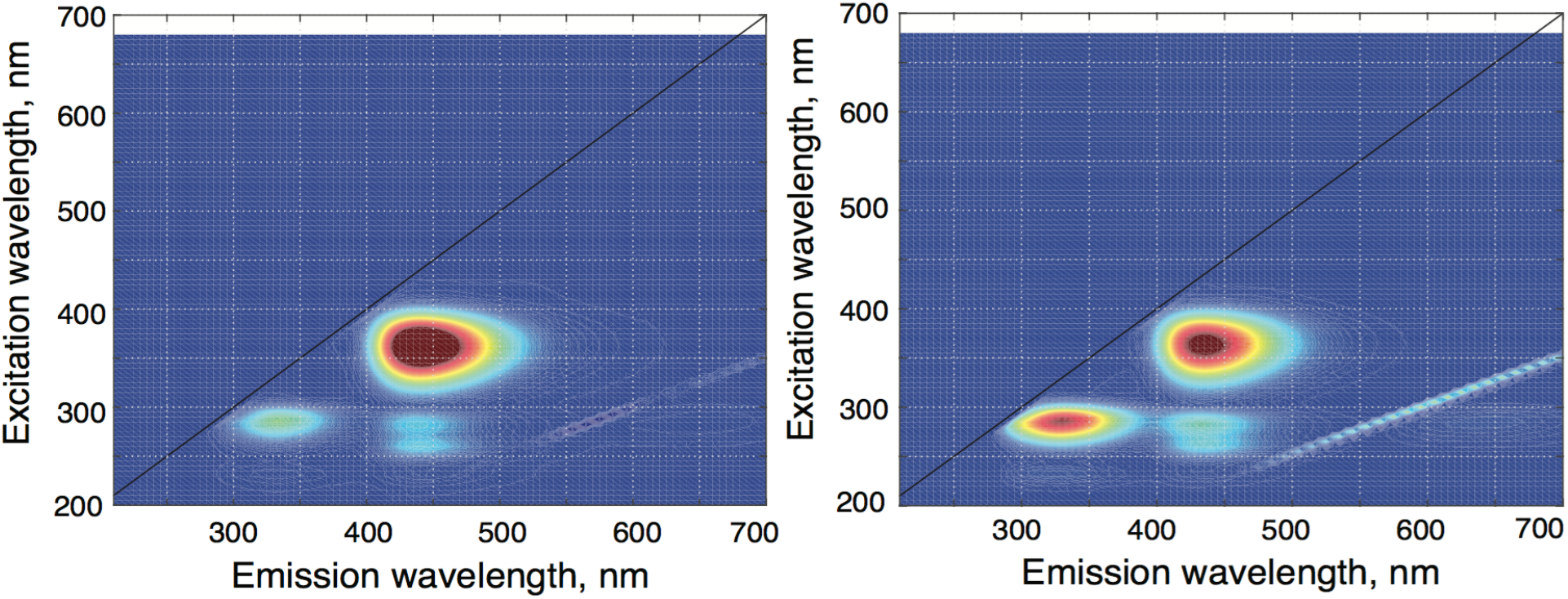
Two-dimensional excitation-emission spectra of the full-length lipidated C118S N-Ras bound with mGDP (left) and mGppNHp (right). The spectra were obtained and processed with *Fluorescence2D*^*104*^.

#### Calculation of the surface density for lipidated peptide added to LUVs assuming even distribution of peptide across LUV surface

Extruder membrane size of 0.2 μm results the radius of LUV around 100 nm;

Therefore:

*LUV surface area* = 4*πr*^2^ = 12.6’10^4^ nm^2^ = 0.126 μm^2^

*Surface area of the lipid (avg)* = 0.7 nm^2^ was estimated by Israelachvili and Mitchell^105^

*Number of surface lipids per LUV* = *LUV surface area / Surface area of the lipid* = 1.8’10^5^ lipids

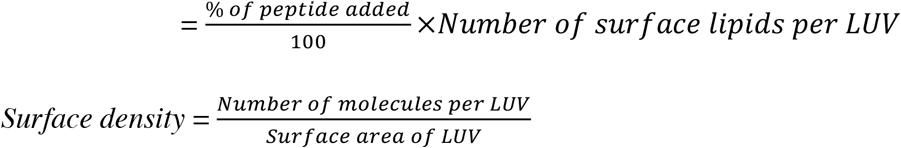

| % of peptide added | Number of molecules per LUVs | Surface density, molecules/$\mu\text{m}^2$ |
| --- | --- | --- |
| 0.1% | 180 | 1400 |
| 0.5% | 890 | 7100 |

#### Estimation of number of lipid rafts per LUV

Size of the lipid raft is temperature-dependent. Pathak and London estimated that size of lipid raft changes form ~150 Å at 10°C to less than 40 Å at 45°C^*50*^. For this calculation, we used average radius of 100 Å (10 nm). Assuming that the raft shape is circular:

*Area of lipid raft* = *π·ŕ* = *π*\*(10)^2^ = 3.14’10^−16^ m^2^

Maximum possible number of rafts per LUV = LUV surface area / Area of raft = 400; Rafts are estimated to occupy approximately 10-40% of the membrane^*106–108*^ in the cell and 50% in the lipid mixture that we are using^*50*^, which results in ca. 200 rafts per LUV. Since the egg sphingomyelin is more hydrophobic than brain sphingomyelin used by Pathak and London^*50*^, it is expected to make larger rafts and this number will be an upper estimate.

#### Estimation of the length of lipid raft boundary and its occupancy with peptide in case it is boundary-associated

Assuming that lipid raft has a circular shape, the length of the boundary for one lipid raft is *C=2×π×r* =62.8 nm

Using an average number of lipid rafts per LUV is estimated above, we can calculated the total length of the lipid raft boundary per one LUV:

Total raft boundary = 1.3 ’10^*-5*^ m

At 0.1% mol/mol of peptide to the total lipid, we will have 70 nm of boundary per one peptide molecule, and 14 nm at 0.5% mol/mol ratio.

Taking into account that lipidated peptide has 7 amino acids (0.8 nm per one amino acid), the estimated length of the extended peptidic part is ca. 6 nm. In the simplistic estimate, we may assume that while the lipid modificaitons anchor the peptide at the boundary, the peptide chain is extended along the boundary helping shield hydrophobic mismatch of the thickness of the raft and disordered phases from aqueous solvent. Janosi et al. reported simulations of H-Ras lipopeptides at the membrane surface, where Ras peptides average end-to-end distance was about 1.2 nm{Janosi, 2012 #6044}. Since the actual conformation of N-Ras lipopeptide at the raft boundary is unknown, we will use 1 nm to 6 nm as a range and arrive at the 1-9% of boundary occupied at the 0.1% peptide concentration and 7-40% at 0.5%.

#### Calculation of the protein surface density for Ras-LUV complex

To calculate protein surface density, we will relate experimentally measured protein content of LUV sample to the total area of LUV in them. To estimate the outer surface area of LUV, we use quantity of total lipid in solution measured by absorption of Rhod-DOPE. The total number of lipids per LUV in both leaflets is^*109*^

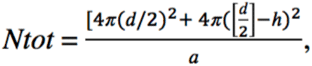

where *d* is a diameter of LUV (known from an extruder membrane pore size), *h* is a thickness of the bilayer (~5nm), *a* is the lipid head group area (*a* for POPC is ~0.71 nm^*2 105*^.

If *d* = 200 nm, then N_tot_ = 3.4 ’10^5^ lipids or 5.7 ’10^−19^ moles; Molar concentration of lipid determined using Rhod-DOPE absorbance in eluted fractions (C_Rhod-DOPE_ = 0.61 μM) gives total lipid concentration considering fraction of Rhod-DOPE added to the lipid mix by experimental design (2%)

*C*_total lipid_ = 31 μM;

Number of LUV in the sample (per L):

*Nlipo* = 5.4 ’10^13^ LUV/L

With this result, the external surface area of LUV per liter of the sample is 6.8 m^2^.

Concentration lipidated Ras molecules in Ras-LUV samples obtained by size exclusion chromatography was estimated from Bradford assay:

| | Concentration<br>measured, $\mu\text{g/ml}$ | Concentration in mol/L |
| --- | --- | --- |
| homogeneous LUV sample | $9 \pm 2$ | $4.5 \times 10^{-7} \text{ M}$ |
| raft LUV sample | $3 \pm 2$ | $1.5 \times 10^{-7} \text{ M}$ |

Calculating molar surface density of Ras on LUV and converting it to units of molecules per square micrometer gives 40,000 μm^2^ for homogeneous and 13,000 μm^2^ for raft LUV samples.

